# A developmental cell atlas of the human thyroid gland

**DOI:** 10.1101/2024.08.22.609152

**Authors:** Hassan Massalha, Mi K. Trinh, Erick Armingol, Liz Tuck, Alexander Predeus, Pavel Mazin, Carmen Sancho-Serra, Agnes Oszlanczi, Yvette Wood, Conor Parks, Toochi Ogbonnah, Holly J. Whitfield, Iva Kelava, Sam Behjati, Roser Vento-Tormo, Nadia Schoenmakers

**Affiliations:** Wellcome Sanger Institute, Hinxton, UK; University of Cambridge, Cambridge, UK; Cambridge University Hospitals NHS Foundation Trust, Cambridge, UK; Department of Paediatrics, University of Cambridge, Cambridge, UK; Department of Metabolism & Systems Science, School of Medical Sciences, College of Medicine and Health, University of Birmingham, UK; Institute of Metabolic Science, University of Cambridge, UK

**Keywords:** Thyroid development, Thyrocyte, Trisomy 21, Thyroid cancer, Single cell genomics, Spatial transcriptomics

## Abstract

The primary function of the thyroid gland is the synthesis and release of thyroid hormones, which are essential for health from embryogenesis to adulthood. Thyroid disorders occur frequently and include congenital hypothyroidism, which occurs due to aberrant thyroid development (thyroid dysgenesis) or impaired hormone synthesis and is particularly prevalent in trisomy 21 (T21). In contrast, thyroid carcinoma, an acquired disorder, is the most common endocrine malignancy in both paediatric and adult populations. Understanding the molecular basis of thyroid dysgenesis and paediatric thyroid carcinoma remains challenging, and requires an improved understanding of foetal thyroid development. To address this, we generated a comprehensive spatiotemporal atlas of the human thyroid during the first and second trimesters of pregnancy. Profiling over 200,000 cells with single-cell sequencing revealed key cell types involved in thyroid gland development, including the hormone-producing thyrocytes. We discovered that foetal thyroid follicular cells are heterogeneous epithelial populations consisting of two main functional subtypes (fTFC1, fTFC2), with fTFC2 expressing increased levels of *PAX8*, and spatial transcriptomics revealed subtype co-occurrence within individual follicles. While both fTFC1 and fTFC2 persist in adult thyroid, fTFC2 is a minor population amongst additional *PAX8*-positive follicular cell subsets. We observed thyroid dysgenesis in T21 age-matched specimens, and T21 thyrocytes showed transcriptional signatures of cytoskeletal disorganisation and altered interactions with the extracellular matrix, as well as compensatory activation of metabolic stress gene programs and upregulation of thyroid biosynthetic genes. In line with the altered proportions of fTFC2 in healthy foetal and adult thyroid, papillary thyroid cancer in children is transcriptionally enriched for the fTFC2 signature compared to that in adults. All together, these findings reveal thyrocyte heterogeneity across the lifespan and provide insights into thyroid development in health and disease, informing potential therapeutic interventions.

## Introduction

The thyroid gland synthesises the thyroid hormones (TH) thyroxine and triiodothyronine, which regulate metabolism, growth and development^1^. It has a bilobed structure composed predominantly of thyroid follicular cells, with a small population of parafollicular cells, also known as C-cells, within a stroma of immune cells, fibroblasts and vasculature. The parathyroid glands are endocrinologically distinct structures located in close proximity to the thyroid that synthesise parathyroid hormone.

Thyroid organogenesis begins at 3 post-conception weeks (PCW) with the specification and proliferation of endodermal precursors and subsequent thyroid budding. By 5.5-7 PCW, the thyroid migrates to its pretracheal position^2–4^, and around 9 PCW folliculogenesis starts, which involves epithelial cell polarisation^5–7^ and lumen formation^5^. Differentiated thyroid follicular cells and C-cells begin hormone synthesis in the second trimester^7^ but continue maturation postnatally. Thyroid development and hormonogenesis requires the coordinated expression of specific transcription factors including *NKX2-1, PAX8, FOXE1* and *HHEX*^8^, followed by upregulation of components of the TH biosynthetic pathway including iodide transporters (e.g. *SLC5A5*, *SLC26A4*), enzymes (e.g. *DUOX2*, *TPO*), and thyroglobulin (*TG*), the scaffold protein on which thyroid hormone synthesis occurs^1,2^.

Aberrant thyroid development and growth leads to neonatal and paediatric thyroid disease. Primary congenital hypothyroidism, the commonest neonatal endocrine disorder (incidence approximately 1 in 2500 neonates), most frequently occurs due to failure of normal thyroid development resulting in athyreosis, thyroid ectopy, or hypoplasia^9,10^. Congenital hypothyroidism, which can be associated with mild thyroid hypoplasia, smaller follicles and mild foetal hypothyroidism^11,12^, also occurs frequently in the context of the genetic disorder Trisomy 21 (T21, also called Down’s syndrome). Furthermore, it has been proposed that aberrant development may play a critical role in the pathogenesis of cancers arising in children - a key distinction from those in adults^13^. Indeed, the transcriptomes of many childhood cancers closely resemble foetal cells, whereas adult tumours are more similar to postnatal cells^14^. Paediatric thyroid cancer, although much rarer than its adult counterpart, remains the most common endocrine malignancy in children^15,16^. The treatment and management strategies for childhood thyroid cancer have generally been extrapolated from studies of adult neoplasms^16,17^. However, thyroid cancer in children typically presents with more aggressive clinical features, including higher rates of metastases and recurrence, and yet often with a better prognosis^18–20^. Moreover, the driver-mutation landscape in paediatric thyroid cancer is distinct from that in adults^21,22^. Despite their significant health impacts, the molecular mechanisms underlying thyroid dysgenesis, including the link between T21 and congenital hypothyroidism, and paediatric thyroid carcinoma remain incompletely understood.

Recent advancements in single-cell multiomics profiling technologies have enabled the characterisation of human development at unprecedented resolution, generating comprehensive and detailed references essential for disease studies^23^. While the majority of thyroid single-cell transcriptomics studies have focused on adult thyroid in humans^24^ and non-human models (zebrafish^25^, mice^24,26^), the foetal thyroid has recently been profiled^27^. However, only two developmental stages were included in this study, and the spatial distribution of cells in tissues remains unknown. Additionally, while several cellular atlases of adult thyroid carcinoma have been generated, research on childhood thyroid cancer remains limited and restricted to bulk RNA sequencing data^28^. The lack of a comprehensive foetal reference has hindered the ability to explore the transcriptional imprints of “foetalness” in thyroid cancer arising in children and adults. Similarly, cell atlases of thyroid in the context of T21 have not been performed, with conclusions primarily drawn from histological analyses. Therefore, there is a significant unmet need to elucidate the cellular and molecular mechanisms underlying thyroid development and its associated pathologies.

In this study, we generated the most comprehensive spatiotemporal atlas of the developing human thyroid. Using this as a reference, we examined the effects of T21 on thyroid cell composition and tissue organisation during development. Finally, we measured foetal cell signals in transcriptomes of thyroid cancer arising in children and adults.

## Results

### An atlas of human thyroid development

To build an atlas of human thyroid development, we performed single cell mRNA sequencing (scRNA-seq) to define the constituent cell types of 27 foetal thyroid glands (9 to 20 PCW) **(Figure 1a, Supplementary Table 1)**, and mapped these cell types onto 4 foetal thyroid glands (12-16 PCW) using spatial transcriptomics (Visium) and multiplexed single-molecule fluorescent in situ hybridization (smFISH) **(Figure 1a)**. We categorised samples into three age windows corresponding to key stages of thyroid development: 9-10 PCW, differentiated thyrocytes; 11-13 PCW, developing follicles; and 14-20 PCW, maturing follicles **(Figure 1b)**.

**Figure 1.**
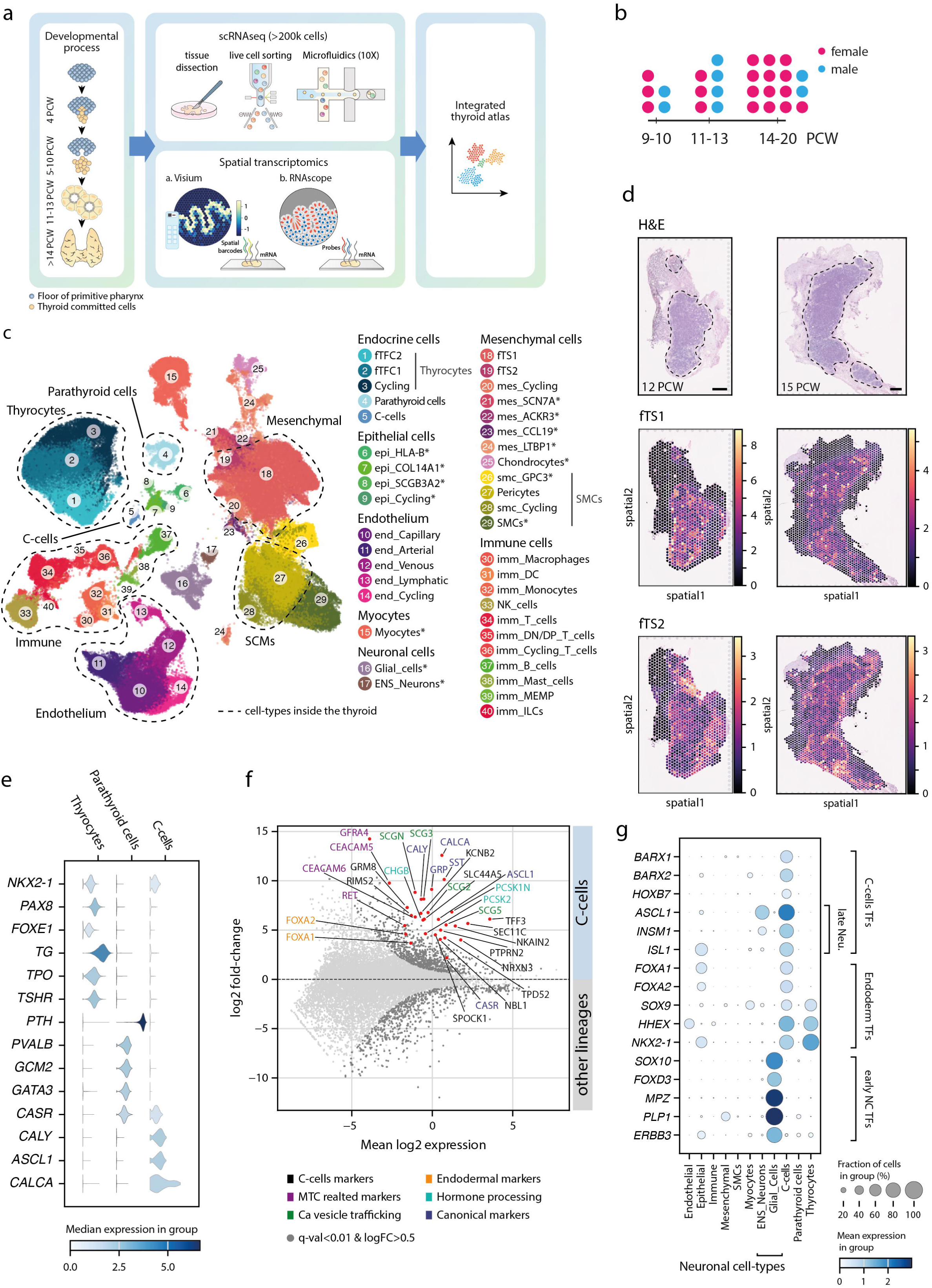
Developmental human foetal thyroid atlas. **a)** Schematic representation of sample collections and workflow. (left-to-right) samples were collected and subjected to both scRNA-seq and spatial transcriptomics analyses, followed by data integration and analysis. **b)** Summary of the 2n samples collected, categorised into three age groups corresponding to key stages of thyroid development: 9-10 (differentiated thyrocytes), 11-13 (developing follicles), and 14-20 (maturing follicles) PCW. Sample sex is indicated by colour. **c)** Uniform manifold approximation projection (UMAP) visualisation of the scRNA-seq data integrated across all collected samples in (b), coloured by cell types. Dashed lines mark cell types present within the thyroids. Asterisks indicate potential contaminant cell types from neighbouring tissues. Abbreviations: epi - epithelial, end - endothelial, smc - smooth muscle cells, mes - mesenchymal, imm - immune, ENS_Neurouns - Enteric neurons, fTFC1 - foetal thyroid follicular cell 1, fTFC2 - foetal thyroid follicular cell 2, fTS1 - foetal thyroid stromal cell 1, fTS2 - foetal thyroid stromal cell 2. **d)** H&E tissue staining of representative samples used in Visium analysis (n=2 biological replicated per analysed age-group). (top left) Dashed line outlines the thyroid gland boundary. Consecutive panels show cellular abundance estimated by cell2location for foetal thyroid stromal cells (fTS1 and fTS2). Sample age indicated in the top panel. Scale bar is 500µm. **e)** Violin plots representing median expression of canonical markers for endocrine cells (thyrocytes, parathyroid cells, and C-cells) captured in the thyroid gland. **f)** M-A plot showing the log2 fold change of gene expression level in the developing C-cells compared to all other cell types (y-axis), and the corresponding average normalised expression level (x-axis). Significantly differentially expressed genes (q-val<0.01 and log2FC>0.5) are shown in dark grey dots. Red dots denote genes previously described in bulk transcriptomic studies and selected top expressed genes. Gene labels are coloured by expression context. Full list of differentially expressed genes is provided in **Supplementary Table 2**. **g)** Dot plot showing mean expression level of canonical transcription factors (TFs) of C-cells, endoderm TFs, and early neural crest (NC) TFs across cell lineages. Transcription factors known to be expressed by differentiated neurons are marked as “late Neu”.

Our single-cell profiling of 216,250 high-quality cells unveiled the cellular composition of the developing thyroid gland **(Figure 1c)**. Cell annotation was performed based on the expression of established markers for different cell lineages^27^ **(Supplementary Figure 1a-b, see Methods)**. In addition to the three main endocrine lineages (thyroid follicular cells, parathyroid cells and C-cells), we found endothelial cells, mesenchymal and immune cells **(Supplementary Figure 1c-d)**. To explore the spatial distribution of the various mesenchymal cell types across the thyroid tissue, we employed the cell2location^29^ algorithm to deconvolute the Visium voxels. Two out of 7 mesenchymal cell types (foetal thyroid stroma 1, fTS1; foetal thyroid stroma 2, fTS2) were located inside the thyroid tissue **(Figure 1d)**. The remaining mesenchymal subsets were found outside the thyroid **(Supplementary Figure 2 and 3),** and along with myocytes, neuronal cells and early epithelial cells are likely contaminants from surrounding tissues. Although all cell types were present across every foetal age profiled, we observed a significant increase in the proportion of fTS1 and pericytes after 10 PCW (p-value < 10^-16^, one-tailed two-proportion z-test), aligned with folliculogenesis and thyroid maturation stages **(Supplementary Figure 1e)**.

We transcriptomically characterised endocrine cell types within the developing thyroid. Thyrocytes expressed key markers *NKX2-1, PAX8, FOXE1*, *TG*, *TPO* and *TSHR* **(Figure 1e)**. Parathyroid cells were also captured due to their close anatomical proximity to the thyroid, and expressed the canonical markers *PTH*, *PVALB*, *GCM2*, *GATA3*, and *CASR*. *CASR*, which aids in monitoring calcium concentration, is also expressed in C-cells **(Figure 1e)**. C-cells are a rare cell population within the foetal and adult thyroid gland, accounting for 270 cells in our dataset. They exclusively expressed *CALY*, *CALCA* and *ASCL1*, with *ASCL1* previously known for its role in the differentiation of serotonergic enteric neurons and C-cells in mice^30^ **(Figure 1e)**. Compared to all other lineages, C-cells also upregulated *GFRA4*^31^, the oncogene *RET*^32^, the hormone Gastrin Releasing Peptide *(GRP)*^33^, and the neuropeptide Somatostatin *(SST)*^34^, all reported in the in the context of C-cells malignancy. **(Figure 1f, Supplementary Table 2)**. Novel genes upregulated in C-cells include *TFF3, NKAIN2, SEC11C, CSG3, CSGN* **(Figure 1f, Supplementary Figure 1f)**.

Although C-cells were previously thought to derive from neural crest cells, recent lineage tracing studies in vertebrates^35^ including mice^36,37^ have demonstrated their endodermal origin. In keeping with an endodermal lineage in humans, *FOXA1* and *FOXA2* are among the most highly expressed and active transcription factors in C-cells **(Figure 1f-g, Supplementary Figure 1g)**. Additionally, C-cells express other classical endodermal transcription factors (*SOX9, HHEX, NKX2-1*), but not classical neural crest transcription factors (*SOX10, FOXD3, MPZ, PLP1, ERBB3*)^38–40^ **(Figure 1g)**. Furthermore, C-cells express genes encoding for hormone-processing proteins (*CHGB*, *PCSK1N* and *PCSK2)*, which are also found in other endodermally-derived endocrine organs^41^ **(Figure 1f)**. These findings support the hypothesis that human C-cells have an endodermal origin.

Taken together, we have generated a single-cell and spatial transcriptomics atlas of the human developing thyroid, and identified the diversity of endocrine cells and surrounding stromal cells.

### Two thyroid follicular cell states coexist during foetal development

Among the thyroid follicular cells population, we identified two cell states that we named foetal thyroid follicular cell 1 (fTFC1) and 2 (fTFC2), along with an additional cluster of cycling thyroid follicular cells **(Figure 2a)**. All thyroid follicular cell states are present across developmental stages **(Supplementary Figure 4a)**. With advancing foetal age, we observed a decrease in the proportion of the cycling population (from 31% to 13% of all thyrocytes) and a corresponding increase in fTFC2 (from 13% to 27%), while the proportion of fTFC1 remained relatively stable **(Figure 2b, Supplementary Figure 4a-b).** Both fTFC1 and fTFC2 increased their metabolic activity during follicle development (11-13 PCW), as shown by an increase in thyroid metabolic score, which was calculated based on the expression levels of TH core metabolic genes **(Supplementary Figure 4c, Supplementary Table 3)**.

**Figure 2.**
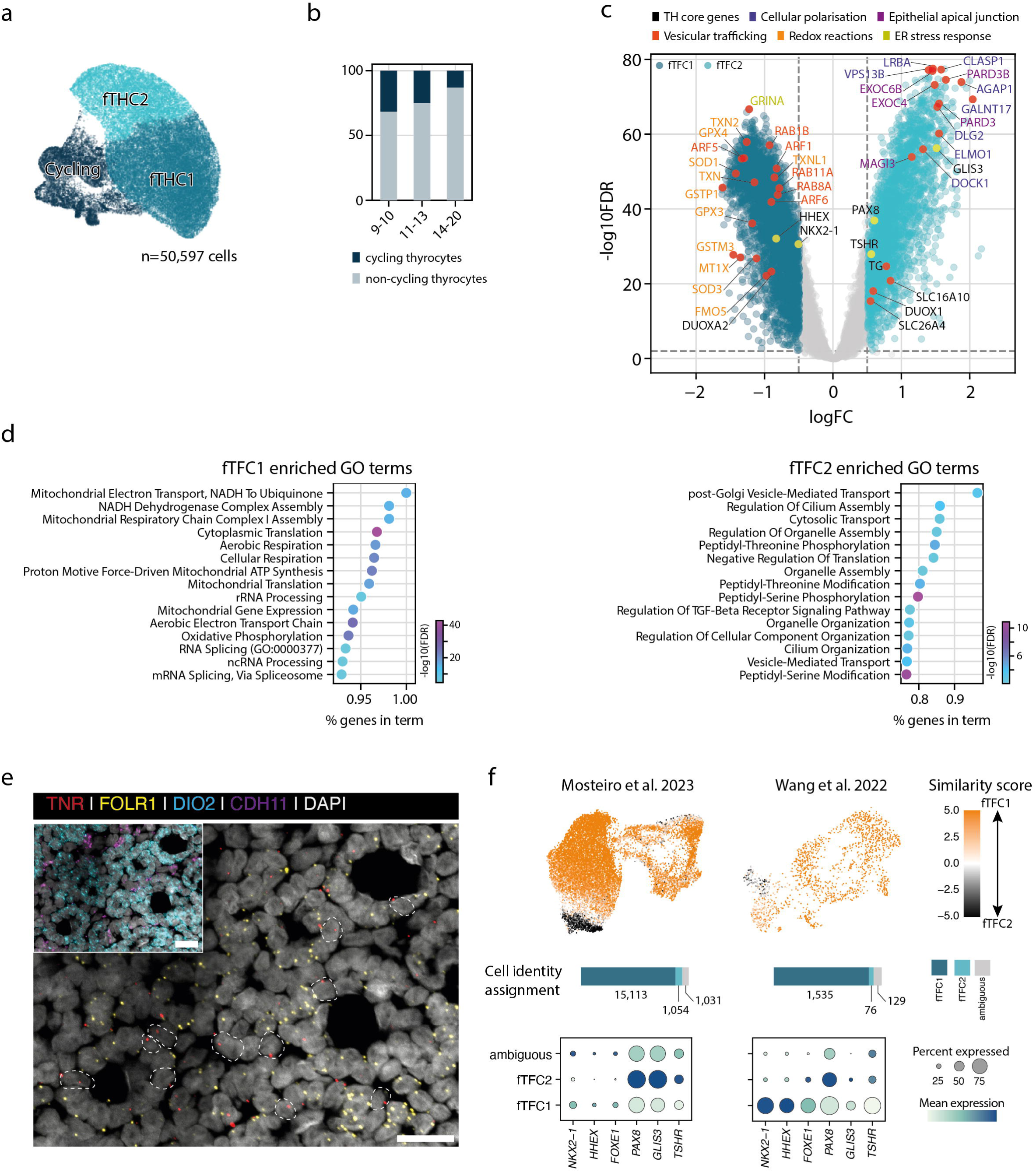
Two subsets of thyroid follicular cells. **a)** UMAP visualisation of thyrocytes coloured by cell-states. ‘n’ is the total number of QC passed cells. **b)** Bar plot showing the proportion of cycling cells among all thyrocytes across developmental age groups. **c)** Volcano plot illustrating the results of the differential gene expression analysis comparing fTFC2 against fTFC1. Coloured dots indicating significant genes (q-val<0.01). Yellow dots show significant TH transcription factors and *TSHR*, red dots indicate annotated genes. Gene labels are coloured by their biological function. **d)** Functional enrichment scores for Gene Ontology (GO) terms calculated using the hypergeometric test for fTFC1 (left) and fTFC2 (right). Terms are sorted by the percentage of genes in each term and coloured by the corresponding FDR. **e)** Spatial localisation of fTFC1 (labelled by *FOLR1*) and fTFC2 (labelled by *TNR*) using RNAscope imaging. The inset image provides an overview of the analysed section stained with *DIO2* (a general thyrocyte marker) and *CDH11* (a general mesenchymal marker). Nuclei are labelled by DAPI (shown in white). A representative image from two independent replicates at 15 PCW is shown. Dashed lines highlight *TNR* positive cells within selected follicles. Scale bar is 20µm. **f)** Top – UMAP of adult normal thyrocytes, extracted from two published scRNA-seq datasets. Cells (dots) are coloured by similarity scores against the reference fTFC1 and fTFC2 cell states as predicted by a logistic regression model. Orange indicates more similar to fTFC1; black indicates more similar to fTFC2. Middle – Bar plots showing the proportion and number of adult thyrocytes being assigned as fTFC1-like (dark blue) or fTFC2-like (light blue) based on the similarity score. Cells with a similarity score between −1.5 and 1.5 are considered ambiguous (grey). Bottom – Dot plots showing the average z-scaled expression level of key transcription factors (*NKX2-1*, *HHEX*, *FOXE1*, *PAX8, GLIS3*) and *TSHR* in different adult thyrocyte populations. Dot size represents the proportion of cells within each category with positive expression. Dots are coloured by the mean expression level across all cells in each group, darker blue indicates higher mean expression.

In comparison to fTFC2, fTFC1 cells exhibited distinct metabolic characteristics, marked by the upregulation of gene ontology terms related to mitochondrial activity and RNA processing **(Figure 2c-d, Supplementary Table 4)** and an increased expression of ribosomal genes **(Supplementary Figure 4d)**. Additionally, there was upregulation of genes associated with reactive oxygen species (ROS) regulation (including *SOD1, SOD3, GPX4, GPX3, GSTP1, GSTM3, TXN, TXN2*), genes involved in the negative regulation of endoplasmic reticulum stress response (*GRINA*), and vesicle trafficking proteins (*ARF1, ARF5, ARF6, RAB1B, RAB11A, RAB8A)* **(Figure 2c)**. While both follicular cell states express key thyroid transcription factors and genes essential for hormonogenesis **(Supplementary Figure 4e)**, fTFC2 showed a significant increase in the expression of transcription factors *PAX8* and *GLIS3*, as well as *TSHR*, which mediates TSH-stimulated thyroid growth and hormonogenesis^42^ **(Figure 2c, Supplementary Figure 4e, Supplementary Table 4)**. Additionally, gene ontology analysis revealed fTFC2 upregulated markers of cilium assembly and targeted vesicle transport processes, which are associated with active cell polarisation, as well as pathways involved in post-translational modification of proteins **(Figure 2d)**. Further upregulated genes included those encoding epithelial apical junctions (*PARD3B, PARD3, MAGI3*), apical vesicle trafficking genes (*EXOC6B, EXOC4*), and genes associated with cellular polarisation (*LRBA, VPS13B, CLASP1, AGAP1, GALNT17, ELMO1, DOCK1, DLG2, DLG5*)^43,44^. *NOTCH2* and *JAG1* were also enriched in fTFC2 **(Supplementary Figure 4f)**.

To investigate whether fTFC1 and fTFC2 cell states were anatomically segregated, we performed spatial transcriptomics and imaging analysis using smFISH with RNAscope. SmFISH revealed that fTFC1 and fTFC2 co-existed within the same follicle, with *FOLR1* (specific to fTFC1) and *TNR* (specific to fTFC2) expressed in adjacent cells **(Figure 2e, Supplementary Figure 4g, Supplementary Table 5)**. To further explore the spatial distribution of fTFC1 and fTFC2, we selected the most dissimilar cells at the opposite coordinates of the Uniform manifold approximation and projection (UMAP) for deconvolution of the Visium data using cell2location **(Supplementary Figure 4h)**. This further confirmed the coexistence of both cell states in the same follicle **(Supplementary Figure 4i)** and validated their presence in almost all follicles in the foetal thyroid gland.

We next investigated if similar populations to fTFC1 and fTFC2 exist in the adult thyroid by exploring two published adult thyroid scRNA-seq datasets^24,45^. Using logistic regression analysis, we directly compared the expression profile of individual adult thyrocytes to each of the two foetal reference populations. This revealed that the majority of adult thyrocytes resemble fTFC1, while only a small number of adult cells (<5%) resemble fTFC2 **(Figure 2f)**. fTFC2-like adult thyrocytes show higher expression of the hormonogenic markers *PAX8, GLIS3* and *TSHR* compared to fTFC1-like cells, as observed in our foetal dataset. The fTFC1-like population exhibits further heterogeneous expression of these markers, indicating additional postnatal changes may occur. For example, the fTFC1-like cells consist of two sub-clusters with varying levels of *PAX8* expression: *PAX8*low and *PAX8*intermediate, as originally described in Mosteiro et al. 2023^24^ **(Supplementary Figure 4j)**. However, we found that *PAX8* expression is highest in the fTFC2-like adult thyrocytes defined here.

In summary, we identified two specialised populations of foetal thyrocytes which are also present in the adult thyroid.

### Thyroid follicular cells in T21 samples

We then used our thyroid foetal atlas as a comparator to investigate the molecular changes in foetal thyroids from individuals with T21. T21 thyroids were significantly smaller (p-value=1.17e-5) than age-matched 2n (euploid) samples, suggesting thyroid hypoplasia **(Supplementary Figure 5a-b)**. We analysed six T21 thyroids at 11, 15 and 17 PCW, and integrated the scRNA-seq data with six age-matched 2n samples **(Figure 3a, Supplementary Figure 5c-d, Supplementary Table 1)**. While we identified the same cell lineages as in the 2n analysis, the relative abundances of clusters within lineage varied between the two karyotypes **(Supplementary Figure 5e-f)**. The same thyrocyte states (fTFC1, fTFC2, and cycling thyrocytes) were present in T21 thyroid glands at proportions similar to those observed in 2n individuals **(Figure 3a, Supplementary Figure 5g)**. However, we found differentially expressed genes between T21 and 2n samples **(Figure 3b).**

**Figure 3.**
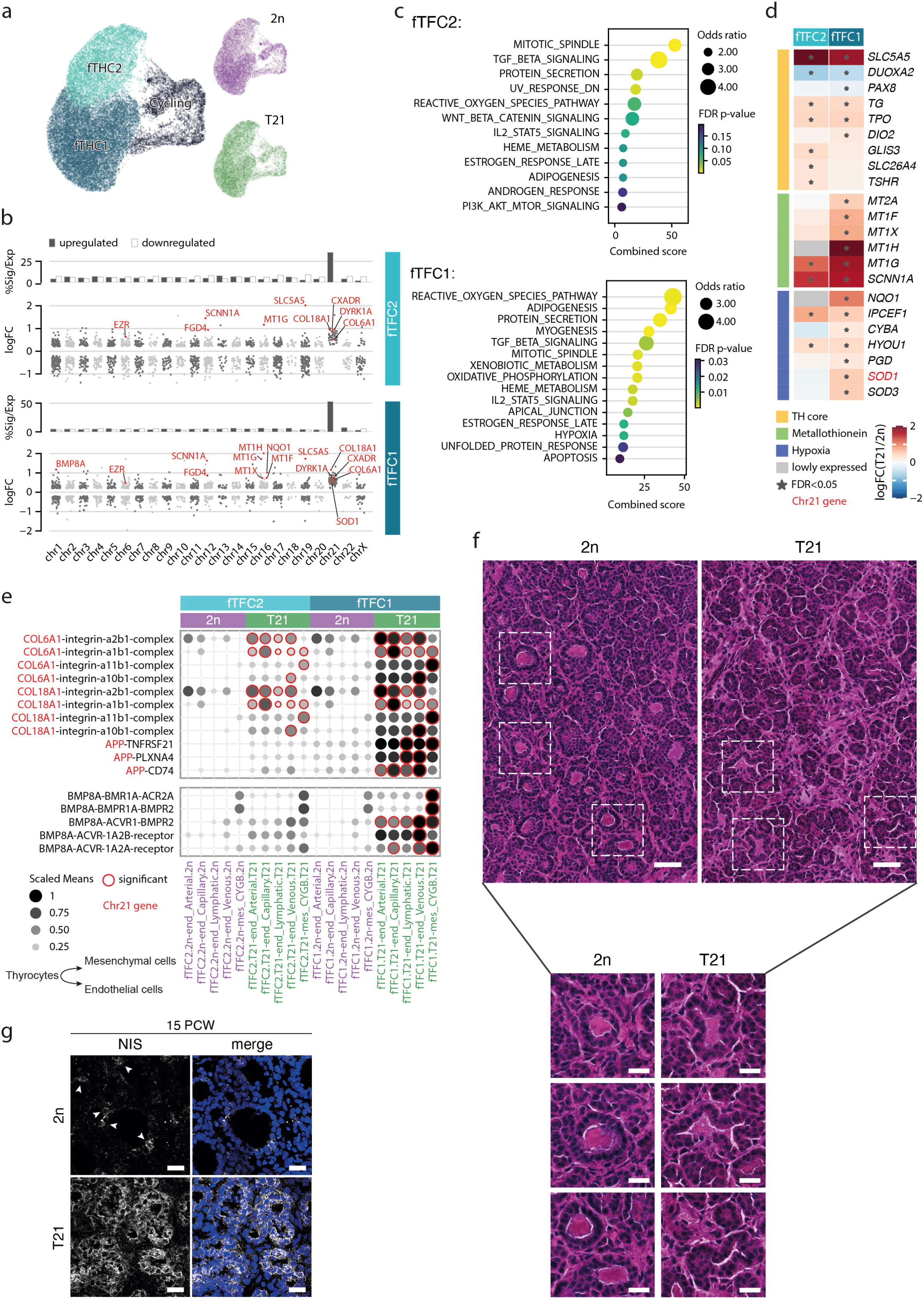
Thyrocytes development in T21 samples. **a)** Integrated UMAP representation of thyrocytes derived from age-matched samples of 2n and T21 karyotypes. Inset panels highlight the distribution of cells from each karyotype within the integrated UMAP. **b)** Significantly differentially expressed genes in T21 compared to 2n for each cell state (FDR<0.05, logFC>0.2), plotted by chromosome. Above each scatter plot, a bar plot shows the percentage of up-/down-regulated differentially expressed genes out of the expressed genes per chromosome. Abbreviations: Sig. - significant, Exp. - expressed. For each cell state, the top significantly differentially up-regulated genes in T21 are highlighted in red. **c)** Gene set enrichment analysis of Hallmark terms for differentially expressed genes per cell state. **d)** Heatmap showing logFC(T21/2n) of selected top differentially expressed genes, as denoted by asterisks and highlighted in (b). Genes are grouped by biological programs which are supported by enrichment terms analysis in (c). **e)** CellphoneDB interaction scores between thyrocytes and mesenchymal/endothelial cells in the thyroid gland for each karyotype separately. X-labels represent interacting cell types, and Y-labels represent interacting ligand-receptor. Selected interactions are presented (for the full list see **Supplementary Figure 6a**). Genes located on chr21 are coloured in red. **f)** H&E staining of thyroid sections prepared from 2n and T21 at 15 PCW. Inset are zoom-in follicles from both karyotypes. Scale bar is 50µm. Inset scale bar is 20µm. **g)** Immunofluorescent images for NIS (encoded by *SLC5A5*) in 2n and T21 thyroids. NIS shown in grey, nuclei labelled by DAPI and shown in blue in merge panels. NIS staining in gray, DAPI counterImages are representative sections from 15 PCW samples. Scale bar is 20µm. Arrowheads indicate the few NIS positive cells per follicle in 2n.

fTFC1 and fTFC2 showed similar numbers of upregulated genes with 9.7% and 6.8% of them located on chromosome 21 (chr21), respectively **(Figure 3b, Supplementary Table 6)**. Gene set enrichment analysis revealed upregulation of genes involved in ‘mitotic spindle’ processes (including: *SPTBN1*, *FGD6, FGD4, EZR, CLIP1, CDC27*) in both fTFC1 and fTCF2, suggesting cytoskeletal alterations in both follicular cell states. Genes implicated in apical junction processes (e.g. chr21 gene *CXADR*), and genes associated with cell polarity (*FGD4*, a direct activator of the essential apical polarity protein *CDC42*^46^ which mediates thyroid bud morphogenesis in mice^47^) were also upregulated in fTFC1. **(Figure 3b,c, Supplementary Table 6)**. Gene ontology analysis also revealed an upregulation of the TGF-beta signalling pathway in both fTFC1 and fTFC2 **(Figure 3c)**, including upregulation of genes such as *BMPR1A, BMPR2, SMURF1*, and *FURIN* **(Supplementary Figure 5h)**. Furthermore, *BMP8A*, a member of the TGF-beta family, was also upregulated specifically in fTFC1 **(Figure 3b)**.

We also observed increased expression of key genes involved in TH biosynthesis (*DIO2, TPO, TG, TSHR*) and iodide transport (*SLC5A5, SLC26A4*) in T21 samples, resulting in an increase in metabolic score **(Supplementary Figure 5i).** In fTFC1 cells there is concomitant upregulation of reactive oxygen species pathway genes *(NQO1, CYBA, SOD1, SOD3, HYOU1, IPCEF1)* and metallothioneins (*MT2A, MT1F, MT1X, MT1H, MT1G*) which are known to protect against ionic stress^48^, as well as the epithelial electrolyte transporter *(SCNN1A)*, which is important for reducing ionic stress^49^ **(Figure 3d)**. The thyrocyte transcriptomic signatures in T21 show upregulation of *GLIS3*, albeit with a small log2FC, whereas other transcription factors critical for thyroid function such as *HEXX, FOXE1, NKX2-1* were not differentially expressed compared to 2n. *PAX8* is slightly downregulated **(Figure 3d)**.

Next, we sought to explore the effect of chr21 triplication on the interaction between thyrocytes and the surrounding stromal compartment by performing CellphoneDB^50^ analysis. Utilising T21 genes differentially expressed by each of the thyrocyte states, we prioritised interactions between thyrocyte-mesenchymal and thyrocyte-endothelial cells in T21 compared to 2n conditions **(Supplementary Figure 6a)**. Notably, the majority of these interactions involved integrins from two main subtypes, arginylglycylaspartic acid (RGD) and collagen receptors, which enhance cell adhesion with the extracellular matrix, promote vascularization, and facilitate collagen interactions^51^ **(Supplementary Figure 6b)**. Among the upregulated ligands expressed by thyrocytes were *COL6A1, COL18A1,* and *APP*, all located on chr21 **(Figure 3e)**. Among the interactions not involving integrins, *BMP8A* signalling was highlighted **(Figure 3e)**. BMP signalling was reported to be crucial for early thyrocytes specification in model organisms^52^.

Histologically, T21 follicles were disorganised, with reduced follicular colloid and smaller invaginated lumens, consistent with previous observations^11^. However, the thyrocyte cellular cytoplasm was expanded, which aligns with the observed increase in metabolic signature. Additionally, we detected an increased amount of extracellular matrix in T21 samples **(Figure 3f, Supplementary Figure 6c)**. Although these histological observations were consistent across all analysed tissue sections, some inter- and intra-individual heterogeneity was noted **(Supplementary Figure 6c)**. Immunofluorescence analysis confirmed the overexpression of NIS protein (encoded by *SLC5A5*) in T21 sections, with elevated NIS levels observed at 15 PCW. NIS, a potent iodine transporter typically expressed in a limited number of cells, showed both increased intensity and a higher number of expressing cells in T21 samples **(Figure 3g)**.

In summary, our findings indicate a significant structural and molecular dysregulation in developing thyrocytes in trisomy 21 samples, which may impact the function of the thyroid gland.

### Thyroid follicular cells in cancer

We next assessed the transcriptional signal of different thyrocyte cell states in thyroid cancer of children and adults. We obtained published bulk transcriptome data of paediatric^53^ (n=37) and adult^54^ papillary thyroid cancer (n=399), the main variant of thyroid cancer **(Supplementary Table 7)**. Using our foetal atlas as reference, we measured signals of different cell types in bulk transcriptomes by performing Cell Signal Analysis as described in Young et.al. 2021^55^. This analysis showed that cell signals in papillary thyroid cancer of both adults and children were predominantly accounted for by foetal thyrocyte signals **(Figure 4a)**. In bulk transcriptomes of normal (non-cancer) adult thyroids, we observed a strong enrichment of fTFC1 signal, with very little or no fTFC2 signal. This is broadly consistent with the aforementioned logistic regression analysis which detected very few fTFC2-like thyrocytes in the adult thyroid **(Figure 2f)**. Similarly, in adult papillary thyroid cancer there was a preponderance of the fTFC1 signal. By contrast, in papillary thyroid cancer of children, we found a significant enrichment of the fTFC2 signal (p-value < 10^-16^, one-tailed Wilcoxon test).

**Figure 4.**
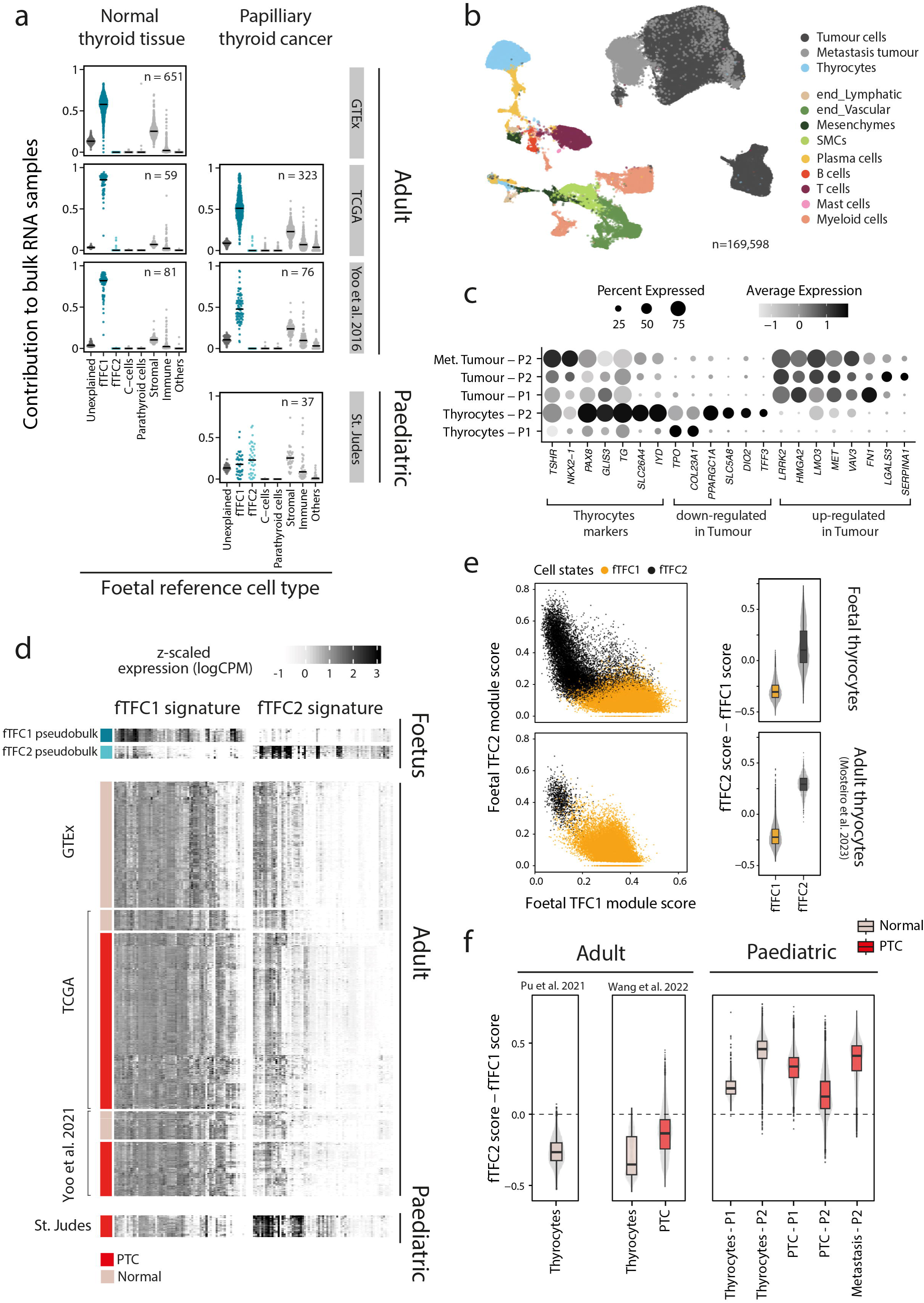
Transcriptional signal of foetal thyrocyte cell states in paediatric and adult papillary thyroid cancer. **a)** The relative contribution of single cell derived signal from the foetal thyroid atlas in explaining the bulk transcriptomes of adult normal thyroid tissues and papillary thyroid cancer in both adults and children. Different foetal reference classes are shown on the x-axis. The relative contribution of each signal to each bulk sample is shown by the y-axis. Each signal - bulk sample combination is represented by a single point. The median of each distribution is shown by the black line. The “n” values denote the number of bulk transcriptomes across different groups. **b)** UMAP visualisation of the paediatric snRNA-seq dataset. Cells (dots) are coloured by their corresponding annotation. Abbreviation: end - endothelial, SMCs - smooth muscle cells. **c)** Dot plot showing the average z-scaled expression level (colour) of canonical thyrocyte and cancer specific marker genes. Dot size represents the proportion of cells within each category with positive expression. **d)** Heatmap showing the z-scaled normalised expression level of fTFC1 and fTFC2 gene signatures in pseudo-bulks of foetal single-cell thyrocytes (top, positive control); bulk transcriptomes of adult normal thyroid tissues and papillary thyroid cancer (middle); and bulk transcriptomes of childhood papillary thyroid cancer (bottom). Z-scaled expression values were obtained by library size normalisation (logCPM), followed by scaling across samples for a given gene. Darker red indicates higher expression level. **e)** Scatter plots showing the enrichment score of fTFC1 signature (x-axis) against fTFC2 signature (y-axis) for individual thyrocytes (dots) extracted from our foetal atlas (top) and the published adult single-cell dataset (bottom). Foetal and adult cells are coloured by the cell state annotation as shown in **Figure 2a** and **2f** respectively. Boxplots showing the corresponding distribution of the difference between fTFC2 and fTFC1 score in individual cells (grey dots) in each cell state (x-axis). Values above zero indicate a higher enrichment for fTFC2 over fTFC1 signature in the given cells. **f)** Boxplots showing the distribution of the difference between fTFC2 and fTFC1 score in individual cells (as in panel e) in normal thyrocytes (grey box) and papillary thyroid cancer cells (red box) in adults (published scRNA-seq datasets) and children (our snRNA-seq dataset).

To extend this observation, we performed single nuclear sequencing of two rare cases of RET-fusion positive papillary thyroid cancers obtained from two girls (pre-schooler; adolescent). We sampled treatment naive resection specimens and collected tissues from tumour, adjacent normal thyroid tissues, as well as a lymph node metastasis in one case **(Supplementary Figure 7a, Supplementary Table 8)**. Following standard pre-processing and quality control steps **(see Methods)**, we recovered expression profiles of 169,598 nuclei **(Figure 4b, Supplementary Figure 7b-e)**. We performed Louvain clustering and UMAP visualisation, which segregated cancer from normal cells, as determined by cell type annotation through label transfer from our single-cell foetal thyroid atlas using CellTypist^56^. In addition, we substantiated the annotation by interrogating the expression of classical papillary thyroid cancer markers across single nuclear transcriptomes **(Figure 4c)**.

To validate the findings of the cell signal analysis in bulk transcriptomes, we derived a list of genes (“gene signature”) that differentiated fTFC1 and fTFC2. The signatures are composed of top differentially expressed genes between fTFC1 and fTFC2 which are efficiently captured in our single-nucleus data **(Supplementary Table 9)**. We then measured the signature in both bulk transcriptomes and single cancer nuclei. We found that the gene signature was sufficient to recapitulate the differential enrichment for fTFC1 and fTFC2 signals in bulk transcriptomes which was detected using the aforementioned analytical approach **(Figure 4d)**. Furthermore, the gene signatures can also differentiate the two thyrocyte cell states in scRNA-seq data, both in foetal and adult dataset **(Figure 4e)**. Assured that the derived gene signatures adequately represent fTFC1 and fTFC2, we quantified the enrichment of their expression in individual normal thyrocytes and papillary thyroid cancer, both in our paediatric single-nuclear dataset and other published adult single cell transcriptomes. We observed that in children, normal thyrocytes and thyroid cancer cells are specifically enriched for the fTFC2 signal, whereas adult counterparts display a stronger fTFC1 signal **(Figure 4f)**. Overall, these analyses indicate that two otherwise indistinguishable neoplasms, adult and childhood papillary thyroid cancer, exhibit different foetal thyrocyte signals on a transcriptomic level.

## Discussion

Despite the frequent occurrence of thyroid dysfunction, thyroid gland development and its cellular heterogeneity in humans are poorly understood. In this study, we presented a comprehensive single-cell and spatial transcriptomics map of the development of the human thyroid gland in 2n and T21 samples. We identified two primary thyroid follicular cell states (fTFC1 and fTFC2) within the developing thyroid gland. These cell states are also found in paediatric and adult thyroids but differ in their proportions, with fTFC2 being more prevalent in younger tissue. We found that T21 profoundly impacts the thyrocyte transcriptome and thyroid follicular architecture, and demonstrated distinct molecular profiles of papillary thyroid cancer in children and adults, with the former enriched for the fTFC2 transcriptional signature.

We used scRNA-seq to analyse 27 euploid thyroid samples collected during the first and second trimester of development, thereby delineating the transcriptome of all cell types within the thyroid. These included hormone-producing thyrocytes and C-cells, as well as parathyroid cells. Although C-cells are a rare cell type, they are clinically significant as the cell population from which medullary thyroid carcinoma originates. Transcriptomic profiling of human C-cells has not previously been undertaken either during foetal development or adulthood. We identify well-validated pathological markers previously derived from bulk studies (e.g., *GFRA4*, *RET*, *CEACAM5*), and other novel markers **(Supplementary Table 2; *CADM1*^57^, *CEP*^58^, *INSM1*^59^, *UCHL1*^60^, *VGF*, *CHGA*, and *CHGB*)**. Future studies are needed to determine their relevance to medullary thyroid carcinoma diagnosis and management. Interestingly, foetal C-cells express somatostatin hormone (*SST*) which has only been previously detected in the context of C-cell malignancy^34^. In keeping with findings from animal models, the transcriptional profile of human C-cells supports an endodermal origin, although in the absence of human lineage tracing studies their provenance would be difficult to confirm.

We identified two foetal thyroid follicular cell states with similarities to the previously characterised *PAX8*low (fTFC1) and *PAX8*hi (fTFC2) subsets in human adult^24,45^, mice^24,26^ and zebrafish^25^. Both cell states express the molecular machinery required for thyroid hormone production. fTFC1 cells are more efficient in ROS scavenging, most likely protecting against the cytotoxic oxidising environment essential for TH biosynthesis (as hydrogen peroxide is required for the oxidation of iodide). In contrast, the fTFC2 gene signature indicates a potential for enhanced hormone synthesis and growth with upregulation of *PAX8*, *GLIS3*, and *TSHR*. Additionally, fTFC2 shows an upregulation in apical vesicle transport processes, which likely supports this enhanced hormone synthesis. Both cell states are detected in normal adult thyroids with similar transcriptomic profiles to the foetal counterparts. However, adult fTFC2-like thyrocytes are much rarer compared to fTFC2 in the foetal thyroid. Moreover, there are two subsets of fTFC1-like cells in adult thyroids: *PAX8*intermediate and *PAX8*low, previously described as thyroid follicular cells 1 and 2^24^. The *PAX8*hi population, corresponding to fTFC2, has not been previously described in adult thyroid.

The specific roles of fTFC1 and fTFC2 and their adult equivalents have not yet been defined. It is plausible that fTFC1 form the main hormone-producing cells, with fTFC2 comprising a ‘reserve’ population capable of augmenting thyroid hormone synthesis or increasing thyrocyte numbers. Similar sensing-related endocrine specialisation has been observed in beta cells^61^. The fact that fTFC2 is present at 9 PCW, prior to folliculogenesis, indicates they could play a crucial role in early development. Further studies are required to understand the plasticity of these subsets, determine whether cells may alternate between them, and characterise their respective functional and proliferative capacities.

Individuals with T21 have a much higher incidence of congenital or childhood-onset mild primary thyroid dysfunction, with mild hypothyroidism also reported during foetal stages^11^. At the population level, T21 is associated with elevated TSH and lower FT4 levels compared to euploid individuals^62^. The most commonly reported thyroid abnormality in T21 is hypoplasia with normal TSH bioactivity suggesting underlying thyroid dysgenesis^63,64^, but the mechanisms underlying this are unclear. Histological analysis of a T21 foetus at 21 PCW revealed small, heterogeneous follicles^11^, consistent with dysgenic thyroid development. Our findings align with this, showing that T21 thyroids are macroscopically hypoplastic, similar to the size reductions observed in other T21 foetal organs, such as the kidney^65^, brain^66^, thymus^67^ and lung^68^. Additionally, T21 thyroids display microstructural disruptions, characterised by smaller follicles, reduced follicular colloid, and irregular lumens.

The T21 thyroid phenotypes likely result from indirect effects of chr21 gene overexpression^69^. Our transcriptomic analysis revealed upregulation of cytoskeletal and apical junction related genes, aligning with the altered follicular morphology. Intriguingly, T21 cerebral organoids^70^ and foetal lung epithelium^68^ show apical polarity alterations, which should be directly investigated in T21 thyroid. T21 thyrocytes exhibit comparable expression of key thyroid transcription factors with euploid thyroid. However, most hormonogenic genes are upregulated, along with enhanced metabolic stress scavenging mechanisms, which may reflect a response to dysregulated hormonogenesis. Furthermore, significant overexpression of *SOD1* and *SOD3* may lead to increased oxidative stress, thereby inducing ROS mitigation pathways.

The increased extracellular matrix deposition we observed in T21 tissue sections is likely partly driven by dose-dependent overexpression of chr21 genes *COL6A1* and *COL18A1*^71,72^, which have also been associated with congenital heart disease in T21^73^, suggesting similar gene dysregulation may be implicated in multiple T21-associated phenotypes. Moreover, given that cardiovascular and thyroid development are coordinated in animal models, and congenital heart disease coexists in 8% of thyroid dysgenesis cases^74^, altered extrinsic signals from mesenchyme and vasculature may also perturb thyroid development^3,47,75^. Consistent with this, interaction analysis predicted elevated TGF-beta signalling (mediated by *BMP8A*) between fTFC1 and mesenchymal/endothelial cells in T21. TGF-beta signalling is known to inhibit thyrocyte proliferation and promote fibrosis^76^, which may explain the increased extracellular matrix deposition in T21 tissues.

Clinically, the thyroid phenotype in T21 is mild and heterogeneous, yet significant at the population level. This aligns with our findings of follicles with normal appearance, and upregulation of components in the TH biosynthesis pathway, particularly NIS, suggests a compensatory mechanism to support hormonogenesis. We also observed inter-individual variability in histological abnormalities, likely influenced by modifier genes, environmental factors, or additional chr21 variation. The transcriptome changes in T21 thyroid share similarities with other organs, with deleterious consequences for thyroid architecture. Further research is necessary to delineate the mechanisms underlying aberrant thyroid development in T21, which may be more widely applicable to understanding the causes of euploid thyroid dysgenesis.

Thyroid cancer in children and adults, apart from variation in the prevalence of specific oncogenic mutations^21,22^, is generally regarded as the same disease, with similar clinical presentations and treatment approaches. In this context, it is remarkable that we were able to detect differences in the enrichment for cell-state-specific transcriptional signals in tumours of adults and children. This may indicate that paediatric and adult thyroid cancers originate from different cell states, or that they assume different cell states upon transformation, perhaps as a consequence of the developmental age of the thyroid gland. This observation was derived from published bulk sequencing data (436 samples), which we further assessed in our independent dataset. Although the single-nucleus data we generated confirms these findings, we would suggest that examining two cases will not have captured the likely cellular variation of paediatric thyroid cancer. To date, there have been very few studies investigating the transcriptomic differences between paediatric and adult thyroid cancers. To enable further mechanistic understandings, systematic international efforts will be required to collect suitable material from these rare cases of paediatric thyroid cancer.

In summary, our thyroid developmental survey is the most comprehensive cellular atlas of the human thyroids to date. This atlas provides new avenues for studying thyroid biology in health and disease and serves as a valuable reference for ongoing advances in *in-vitro* models of thyroid gland development^27,77–79^.

## Supplementary Tables

Supplementary Table 1 - List of foetal samples used in the study

Supplementary Table 2 - Differential gene expression of C-cells

Supplementary Table 3 - Thyroid metabolic score genes

Supplementary Table 4 - Thyrocytes differential gene expression fTFC2 vs fTFC1

Supplementary Table 5 - RNAscope probes used in the study

Supplementary Table 6 - Thyrocytes differential gene expression T21 vs 2n

Supplementary Table 7 - List of publicly available paediatric and adult bulk transcriptomes used in this study

Supplementary Table 8 - List of paediatric samples generated in this study

Supplementary Table 9 - List of genes included in the fTFC1 and fTFC2 signatures

## Supplementary Figure legends

**Supplementary Figure 1**

**a)** UMAP plot displaying identified lineages in the study. ‘n’ is the total number of QC passed cells. **b)** Dot plot illustrating the expression levels of the top 5 markers for each lineage sorted by fold change. Expression values calculated from the full dataset. **c, d)** Distribution of sample ages (c), and karyotypes (d) of the identified cell types in Figure 1c. **e)** Bar plot showing proportion of cell types found within the thyroid gland across a developmental age bin. **f)** Normalised expression of top 30 genes expressed in C-cells. Reported are significant genes (FDR<0.01) and grouped by functional annotations. Presented expression values calculated from the full dataset. **g)** Z-score of C-cells TFs activity calculated using Dorothea^80^.

**Supplementary Figure 2**

Abundance of mesenchymal and SMCs cell types estimated using cell2location for a 12 PCW sample. The first panel on the left displays an H&E stained section adjacent to the Visium analysed section. Presented is a selected sample from two independent replicates. Scale bar is 500µm.

**Supplementary Figure 3**

Abundance of mesenchymal and SMCs cell types estimated using cell2location for a 15 PCW sample. The first panel on the left displays an H&E stained section adjacent to the Visium analysed section. Presented is a selected sample from two independent replicates. Scale bar is 500µm.

**Supplementary Figure 4**

**a)** Composition of thyroid follicular cell states across different age groups. **b)** Composition of thyroid cycling cells across sample age shown in a UMAP (left) and barplot (right). **c)** Thyroid metabolic scores for thyrocytes across all PCWs. **d)** Volcano plot comparing fTFC1 vs fTFC2, highlighting majority of ribosomal genes upregulated in fTFC1. **e)** Line plots showing the mean expression of sum-normalised UMI counts for thyroid TFs and core metabolic genes across age groups for fTFC1 and fTFC2. Dots represent normalised mean expression at each time point. Shaded areas representing ±SEM. **f)** UMAP showing normalised expression levels of *NOTCH2* and *JAG1*. **g)** UMAP showing normalised expression levels of *TNR* and *FOLR1* used for RNAscope imaging. **h)** Edge clusters for fTFC1 and fTFC2. Highlighted cells used for spatial mapping of fTFC1 and fTFC2 via cell2location. **i)** H&E staining alongside abundance analysis of fTFC1 and fTFC2 using cell2location, based on edge Leiden clusters identified within each of the two analysed cell states (g). Sample age is indicated in each panel. Scale bar is 500µm. **j)** UMAP of adult normal thyrocytes, extracted from published scRNA-seq datasets: Mosteiro et al. 2023 (top panels) and Wang et al. 2022 (bottom panels). Cells (dots) are coloured by: (left column) similarity scores against the reference fTFC1 and fTFC2 cell states as predicted by a logistic regression model (as shown in Figure 2f), and (right column) normalised expression level of *PAX8*. Abbreviations: *PAX8*int - *PAX8* intermediate, *PAX8*hi - *PAX8* high.

**Supplementary Figure 5**

**a)** Macroscopic images displaying developmental thyroid gland tissues from 2n and T21 samples at 12 and 15 PCW. The images are captured at the same magnification, highlighting the delayed growth of the T21 thyroid gland. Scale bar is 2mm. **b)** Mean lobe length in [mm] plotted against sample age [PCW] for each karyotype: T21 (green) and 2n (purple). Solid lines represent the linear regression fit for each karyotype, with shaded areas indicating the 95% confidence intervals. A p-value of 1.17e-05 indicates a statistically significant difference in growth patterns between the two karyotypes. Analysis of covariance (ANCOVA) test was performed to compare the two datasets. L1 and L2 represent the maximum lobe lengths measured from the sample midline per sample, as shown in the schematic representation on the bottom-left. **c)** Summary of age-matching samples used for T21 transcriptional analysis. Samples categorised into two age groups: 11 and 15-17 PCW. Sample sex is colour-coded. **d)** Mean normalised expression levels of genes located on chr21. T21 samples exhibit higher mean expression due to the additional copy of chr 21. **e)** Integrated UMAP representation of high-quality cells derived from the samples in (c). Annotations are cluster names. LECs - lymphatic endothelial cells, VECs - vascular endothelial cells, ILCs - Innate lymphoid cells, ENS_Neurouns - Enteric neurons, SMCs - smooth muscle cells. **f)** Karyotypes distribution within cell clusters annotated in (e). **g)** Karyotypes distribution within thyrocytes states. **h)** Scaled mean expression of ‘TGF-beta’ term enriched genes. Groups are cells within each cell state per karyotype. **i)** Thyroid metabolic score calculated per cell state for each karyotype. Significance calculated using Wilcoxon rank-sum test. **** is p-value<=1.0e-4.

**Supplementary Figure 6**

**a)** Complete output of CellphoneDB interactions between thyrocytes-mesenchymal and thyrocytes-endothelial cells. Significant interactions are marked by red rings. Genes located on chr21 are labelled in red. Interacting cell types from T21 samples are labelled in green, and those from 2n samples are labelled in purple. **b)** Summary of CellphoneDB interactions in (a) grouped by thyrocyte states, showing integrins as the dominant interaction partners. Numbers in ring segments [n/m] indicate the interaction count of a given group [n] relative to the total interactions in the parent group [m]. Most interactions involve two main integrin subtypes. **c)** H&E staining of 2n and T21 thyroid sections, depicted in Figure 3f, highlights extracellular regions that are more prominent in T21 samples. Images are from different donors than those shown in Figure 3f. Scale bar is 50 µm.

**Supplementary Figure 7**

**a)** Schematic overview of childhood papillary thyroid cancer sample collection and data generation. Asterisk denotes the driver event in each thyroid cancer case. **b-c)** UMAP visualisation of the paediatric single-nucleus RNA-seq dataset. Cells (dots) are coloured by their corresponding donor (**b**) and sampling site (**c**). **d)** Barplot showing the proportion of cells coming from each donor across cell types. **e)** Dot plot showing the average z-scaled expression level (colour) of canonical and cell type specific marker genes. Dot size represents the proportion of cells within each category with positive expression. Met for metastasis.

## Methods

### Donors samples

Human embryonic thyroid samples was provided by the Joint MRC / Wellcome Trust (Grant #MR/006237/1) Human Developmental Biology Resource (http://www.hdbr.org). Paediatric normal and thyroid cancer samples were collected through studies approved by the National Health Service National Research Ethics Service reference 16/EE/0394. Informed written consent was obtained from all patients or their guardians.

### Tissue processing

Thyroid glands dissected from human embryos of ages 9-20 PCW, washed with ice-cold phosphate buffered saline (PBS) solution (Gibco, 10010023) and immediately transferred either into ice-cold HypoThermosol^®^FRS solution (Sigma-Aldrich, H4416) and kept at 4°C for same day fresh sample processing or into cryopreserved media and stored at −80°C (see below). For large samples, part of the tissue ∼0.3 cm^3^ was embedded in optimal cutting temperature (OCT) compound (ThermoFisher Scientific, 23730571) inside a cryomold and rapidly frozen in dry ice/isopentane slurry for spatial analysis.

### Tissue dissociation for scRNA-seq

In the case of cryopreserved samples, tissues were thawed at 37°C, transferred to a 15 ml tube and topped-up with 13 ml of ice cold 10% RPMI/FBS (Gibco, 1640). Samples were centrifuged (450 x g, 5 min, 4°C). The supernatant was discarded and the samples went through the same digestion steps as for fresh samples. Fresh samples washed once with ice-cold 10% RPMI/FBS, chopped with a sharp scalpel and transferred into 15 ml tube containing per-warmed enzymatic digestion mix (10% RPMI/FBS containing Collagenase IA (Sigma-Aldrich, C2674), Liberase TM (Roche, 5401119001), and DNAse I (Roche, 11284932001) with final concentrations of 1 mg/ml, 50 ug/ml, and 0.1 mg/ml, respectively) for 30 min on a tube rotator at 37°C. Digested tissue centrifuged (450 x g, 8 min), the supernatant removed and 4 ml of 0.25% (v/v) trypsin-EDTA (Sigma-Aldrich, T3924) and DNAse I (0.1 mg/ml) was added to the cells pellet and any undigested tissue and incubated at 37°C for 10 min a tube rotator. The digression process stopped by doubling suspension volume with fresh 10% RPMI/FBS media. Cell suspension was centrifuged (450 x g, 8 min), supernatant was discarded, and 1 ml of red-blood-cell (RBC) lysis buffer (eBioscience, 00-4300) was added for 10 min at room temperature. After the RBC lysis step, cell suspension passed through a 100 µm cell strainer (pluriSelect, 43-50100-01), the filter washed with 3 ml fresh 10% RPMI/FBS and centrifuged (450 x g, 8 min, 4°C). The supernatant was discarded and replaced with 1 ml of ice-cold Cell Stain Buffer (BioLegend, 420201) and kept on ice. To enrich for live cells, live/dead sorting was applied to all samples, except for Hrv159. Propidium Iodide (PI) fluorescent dye (ThermoFisher Scientific, P1304MP) was added at 1:1000 (v/v) to stain dead cells. Live cells were sorted using SONY M900 into an ice-cold Cell Stain Buffer containing tube. Sorted single-cell suspension kept on ice until it was loaded onto the 10X Chromium chip.

### Tissue cryopreservation

On ice, fresh tissue samples were washed twice with cold, fresh 10% RPMI/FBS in a petri dish. After discarding the washing solution, 1 ml of ice-cold Cryostor solution (CS10) (C2874-Sigma) was added to the tissue. The tissues were then cut into segments >1 mm^3^ using a sharp scalpel. The tissue pieces were collected with a wide-open tip and transferred into cryovials and kept on ice. The petri dish was rinsed with 0.5 ml of ice-cold Cryostor solution (CS10), which was also added to the cryovials. The cryovials were then placed in a CoolCell (Corning) and stored at −80°C overnight, allowing the temperature to decrease by approximately 1°C per minute. The next day, the cryovials were removed from the CoolCell and kept at −80°C until further processing.

### Library preparation and sequencing

Sorted cells were counted and loaded according to the manufacturer’s protocol for the 10X Chromium Single Cell 5′ Kit v.02, to recover between 2,000 and 10,000 cells per reaction. In the case of cell clumps observed during the counting step, cells passed through a 70µm Flowmi filter (SP Bel-Art, 136800070). Filtered samples recounted before proceeding with the 10X reaction preparation. Libraries were sequenced, aiming at a minimum coverage of 50,000 raw reads per cell, on the Illumina Novaseq 6000 system; using the sequencing format; read 1: 28 cycles; i7 index: 10 cycles, i5 index: 10 cycles; read 2: 90 cycles.

Raw reads were mapped and quantified using the STARsolo algorithm. STAR version 2.7.10a_alpha_220818 compiled from source files with the "-msse4.2" flag was used for all samples. Wrapper scripts documented in https://github.com/cellgeni/STARsolo/ were used to auto-detect 10X kit versions, appropriate whitelists, and other relevant sample characteristics. Human reference genome and annotation exactly matching Cell Ranger 2020-A was prepared as described by 10X Genomics: https://support.10xgenomics.com/single-cell-gene-expression/software/release-notes/build#header. For 10X samples, the STARsolo command was optimised to generate the results maximally similar to Cell Ranger (v6). Namely, “--soloUMIdedup 1MM_CR --soloCBmatchWLtype 1MM_multi_Nbase_pseudocounts --soloUMIfiltering MultiGeneUMI_CR --clipAdapterType CellRanger4 --outFilterScoreMin 30” were used to specify UMI collapsing, barcode collapsing, and read clipping algorithms. For paired-end 5’ 10X samples, options “--soloBarcodeMate 1 --clip5pNbases 39 0” were used to clip the adapter and perform paired-end alignment. For cell filtering, the EmptyDrops algorithm employed in Cell Ranger v4 and above was invoked using “--soloCellFilter EmptyDrops_CR” options. Options “--soloFeatures Gene GeneFull Velocyto” were used to generate both exon-only and full length (pre-mRNA) gene counts, as well as RNA velocity output matrices.

For cell hashing experiments (HCA_GLNDrna12943535, HCA_GLNDrna12943536, HCA_GLNDrna11814746, HCA_GLNDrna11814737; see Data availability), Cell Ranger (v7.0.1) was used in “cellranger count” mode. TotalSeq B antibody reference was used to assign cell hashing reads. Cell Ranger transcriptome reference 2020-A was used for gene expression quantification.

To perform donor-based demultiplexing, Souporcell (v2.5) was used in “common variant” mode, using 1000 Genomes reference VCF file filtered to retain the variants with allele frequency of 0.05 or greater; additionally, options “--skip_remap True --no_umi True” were set during the processing. Matched donors were identified using Souporcell auxiliary script shared_samples.py.

### Quality control and preprocessing of scRNA-seq data

After demultiplexing the pooled samples, basic filtering was applied (keeping cells with >1000 genes and genes expressed in >3 cells) before downstream analysis. Ambient noise was removed using SoupX (v1.6.2)^81^ pipeline with the default settings. Next, we calculated a doublets score using Scrublet (v0.2.3)^82^. The integrated manifold of the analysed samples were generated using single-cell Variational Inference (scVI; v0.6.8), with donor as batch factor. All the remaining parameters were kept as default, with n_latent=40, n_layers=1. The scVI low dimensional space was estimated on the top 3,000 most highly variable genes, which were defined using ‘*Seurat v3’* flavour on the raw counts. With the resulting scVI-corrected latent representation of each cell, we estimated the neighbour graph, generated a Uniform Manifold Approximation and Projection (UMAP) visualisation and performed Leiden clustering. The resolution of the clustering was adjusted manually so that all the previously described thyroid cell types were resolved^26,27,83^. To aid marking *low QC* clusters, we calculated the following two scores: (i) ‘High Scrublet Score (High-SS)’ assigned for cells with a ‘scrublet’ score >0.25. (ii) ‘Red Blood Cells Score (RBCS)’ were calculated using the scanpy ‘score_genes’ function on log-normalised counts using the following genes *HBB*, *HBG2*, *HBG1*, *HBA1*, *HBA2*, and *HBD*. High-RBCS was assigned to cells with RBCS > 2. Clusters with >20% cells marked as High-SS or High-RBCS are considered *low QC* clusters. A second round of scVI integration was performed after removing *low QC* clusters. Lineage annotations were assigned for the generated clusters based on well-established marker genes of the different cell types (i.e. thyrocytes, epithelial, neuronal, mesenchymal, immune and endothelial cells). Above steps were also applied to generate a 2n-T21 age-matched integrated atlas.

### Annotation of cell types

We performed a zoom-in re-annotation for each lineage in the integrated scRNA-seq manifold. Cell subsets of each lineage were reintegrated with Harmony (harmonypy package; v0.0.5)^84^ with theta = 0. Before integration, we excluded donors with fewer than 10 cells in total. Next, we removed cells with ‘n_genes’ <1000, ‘n_counts’ <2000 and cells with percent of mitochondrial genes >0.2. With the retained cells, we calculated highly variable genes using scanpy ‘highly_variable_genes’ (flavor=’seurat_v3’, n_top_genes=2000). For thyrocytes, we excluded the following genes before retrieving highly variable genes, (i) 30 stress genes^85^, (ii) mitochondrial genes, (iii) ribosomal genes, and (vi) lowly expressed genes using scanpy ‘filter_genes’ function at ‘min_counts=3’. A ‘n_top_genes=4000’ was selected for thyrocytes. With the generated clusters, we performed a quality control assessment to exclude clusters that are likely driven by technical artefacts (i.e. *low QC* cells or doublets). Briefly, *low QC clusters* are those with: (i) >=20% of cells with High-SS, or (ii) expresses a mixture of bona-fide markers of different lineages, and (iii) do not express any distinctive gene (thus do not represent any independent biological entity). After excluding *low QC* clusters, we conduct a second round Harmony integration using the above parameters, with a new PCA dimensions determined based on an elbow plot after removing *low QC* and doublets cells, if any. Distinctive marker genes per cell type were extracted using Term Frequency - Inverse Document Frequency approach (TF-IDF), as implemented in the SoupX package (v1.5.0). TF-IDF analysis was performed on cells with ‘scrublet’ score below 0.2. Top genes/unique markers were used to annotate the different cell types per lineage. To annotate immune cells in our datasets, we first utilised Celltypist (v1.5.2)^56^, a logistic regression classifier optimised by the stochastic gradient descent algorithm^83^. We used the pretrained model ‘Pan_Fetal_Human’ (v2)^86^ to annotate immune cells in our database. Immune cell annotation was determined based on the ‘majority_voting = True’ parameter. In cases where multiple subcell types were assigned to a single cluster, we opted for higher-level annotation. The annotation process described above was applied to the integrated 2n-T21 atlas. Additionally, we utilised a logistic regression model, trained on the 2n atlas, to assist in annotating the integrated atlas for both karyotypes.

To assess whether there was a significant increase in the proportion of fTS1, fTS2 and pericytes after 10 PCW, the one-tailed two-sample test for equality of proportions was employed (using the R function ‘prop.test’, without Yate’s continuity correction) to compare the proportion of each cell type at both 11-13 PCW and 14-20 PCW against that at 9-10 PCW.

### Lineage DGE analysis

All lineages were randomly downsampled to 270 cells each (matching the number of C-cells). Protein-coding genes expressed in a minimum of 10 cells and above 20% of a cluster’s cells retained for downstream analysis. We used the ‘FindAllMarkers’ function from the Seurat package (v4.0.1) to find lineage specific genes shown in **Supplementary Figure 1b**. C-cells differentially express genes reported in **Supplementary Table 2**. Top C-cells TFs in **Supplementary Figure 1f**.

### C-cells TF analysis

TF activity scores in C-cells were computed using Dorothea (v1.5.0)^80^ package. The Univariate linear model (ULM) method was used with the default parameters for a list of ‘source’ and ‘target’ from all levels. T-test was applied to compare TF activity score in C-cells against the rest. TFs with a ‘activity’ mean change >2 and FDR <0.01 were retained for downstream analysis and visualisation.

### Thyroid metabolic score

To quantify the metabolic activity of thyrocytes, we defined the Thyroid metabolic score based on the expression levels of key thyroid metabolic genes (*DIO2, DUOXA1, DUOXA2, DUOX1, DUOX2, SLC5A5, ANO1, SLC26A4, TSHR, TPO, IYD, TG, SLC16A2, SLC16A10*) **(Supplementary Table 3)**. We calculated the Thyroid metabolic score using the ‘sc.tl.score_genes’ function in scanpy with default settings.

### Differential gene expression analysis of the two thyrocytes populations

To identify differentially expressed genes (DEGs) between the two thyrocytes cell-types, while accounting for donors’ age as a covariate, we used the tool edgeR (v3.32.1)^87^. We first generated pseudobulks for each combination of donors (27 levels) and cell-types (fTFC1, fTFC2), allowing 3 technical replicates per donor per condition, following the tutorial at https://www.sc-best-practices.org/preamble.html^88^. Pseudobulks with fewer than 30 cells were excluded, along with non-coding genes and 33 Y chromosome genes. The resulting pseudobulk data was used to fit a generalised linear model using the ‘glmQLFit’ function. The model formula included the following factors: ‘celltype’ with two levels (fTFC1, fTFC2), ‘age-group’ with five levels (9-10, 11-13, 14-15, 16-17, 20), and without an intercept term. A post-hoc analysis on the coefficients of the ‘celltype’ levels was performed to identify genes which are differentially expressed between the cell types. Here, we excluded lowly-expressed genes by only evaluating genes expressed in at least 20 cells and present in more than 10% of either compared cell-type. We corrected for multiple hypothesis testing by using the Benjamini-Hochberg method and a false discovery rate (FDR)<1%.

### Downstream analyses of thyrocytes differentially expressed genes

After identifying differentially expressed genes, a Gene Ontology (GO) enrichment analysis for cell-type-specific genes was performed using Fisher’s exact test. The package ‘gseapy’ (v1.0.5) was used to obtain terms from ‘GO_Biological_Process_2023’. Terms related to "disease, cancer, infection, carcinoma" were excluded, and only terms with at least 10 significant genes were retained. Terms with FDR <0.01 were considered statistically significant.

### Detection of fTFC1- and fTFC2-like cells in published adult thyroid scRNAseq datasets

Published scRNA-seq dataset of the adult thyroids were obtained from Mosteiro et al. 2023^24^ and Wang et al. 2022^45^. Thyrocytes from normal (non-cancer) samples from each dataset were retained for the analysis.

To compare individual transcriptomes of adult thyrocytes against the two reference foetal thyrocyte cell states, we employed a method based on logistic regression as previously detailed in Young et. al. 2018^89^. Briefly, a logistic regression model with elastic net regularisation (alpha = 0.1) was trained on a single-cell transcriptomic reference, using the ‘cv.glmnet’ function in R. Our reference dataset used in this analysis consists of two foetal thyrocyte subtypes fTFC1 and fTFC2, and Schwann cell precursors (from Kildisiute et.al. 2021^40^) as negative control. This model is then applied to calculate the predicted similarity scores for individual cells from the query dataset (adult thyrocytes) against each of the reference classes (i.e. different cell types in the reference dataset). The final fTFC1/fTFC2 similarity score for each cell was calculated as the difference between its predicted fTFC1 and fTFC2 similarity scores.

### T21 vs 2n analysis

Pseudobulks were generated for fTFC1 and fTFC2 cell-states, as described in the ‘Differential gene expression analysis of the two thyrocytes populations’ section in Methods. We generated 3 technical replicates per donor per condition. Pseudobulks with fewer than 30 cells were excluded, along with non-coding genes and 33 Y chromosome genes. We then fitted a generalised linear model (GLM) using the ‘glmQLFit’ function from edgeR (Y ∼ 0 + age_group + karyotype) for each cell-type separately^88^. From this model, a post-hoc analysis on the coefficients of the ‘karyotype’ levels was performed to identify differentially expressed genes between T21 and 2n. Only genes expressed in at least 20 cells and present in more than 10% of either compared cell groups were considered. We corrected for multiple hypothesis testing by using the Benjamini-Hochberg method and a false discovery rate (FDR) <5%. Gene set enrichment analysis was performed on genes with logFC >0.2 using decoupler (v1.5.0) ^80^ focusing on ‘Hallmarks’ terms. Terms with FDR <0.2 were reported.

### Cell-cell interaction analysis

We used CellphoneDB (v5.1)^50^ with CellphoneDB-database (v5.0) to identify interactions between thyrocytes and endothelial/mesenchymal cells. To focus on interactions associated with karyotype differences, we employed the DEGs-based method of CellphoneDB using thyrocyte DEGs in T21 cells (see previous section ‘T21 vs 2n analysis’). Karyotype information was provided as ‘microenvironments’ to ensure that only interactions between cells of the same karyotype were considered. A stringent filtering criteria was applied to refine DEGs (FDR <0.01, logFC >0.5). Genes present in at least 10% of the compared group were considered for this analysis. The resulting interactions were plotted using the ktplotspy package.

### 10X Genomics Visium library preparation and sequencing

10 µm fresh frozen tissue cryosections were collected onto a Visium slide and the 10X Genomics Visium Spatial Gene Expression protocol was followed (User Guide CG000239 Rev F). H&E stained Visium slides were imaged at 40x on a Hamamatsu Nanozoomer 2.0 HT slide scanner. The tissue was permeabilized for 30 mins, a pre-treatment which was optimised in a separate time course experiment (10X Genomics, User Guide CG000238 RevE). Captured cDNA transcripts were then eluted and dual indexed following the 10X Genomics Visium library preparation protocol. Libraries sequencing done on Illumina NovaSeq 6000 SP flow cell.

### Cell2location

To map cell-types identified by scRNA-seq in Visium data, we used the cell2location (v0.1) method^29^. Firstly, we trained a negative binomial regression model to estimate reference transcriptomic profiles for all the cell types profiled with scRNA-seq on two levels of annotation ‘celltype’ and ‘celltype_edge’ (that contains the edge cells of fTFC1 and fTFC2; **Supplementary Figure 4g**). We excluded very lowly expressed genes using the filtering strategy recommended by Cell2location (cell_count_cutoff=5, cell_percentage_cutoff2=0.03, nonz_mean_cutoff=1.12). Individual 10X samples were considered as a batch, and donor was used as categorical covariate. Training was performed for 250 epochs and reached convergence according to manual inspection. Next, we estimated the abundance of cell-types in the spatial transcriptomics slides using reference transcriptomic profiles of different cell-types. All slides were analysed jointly. Following cell2location hyperparameters were used: (1) expected cell abundance (N_cells_per_location) = 45; (2) regularisation strength of detection efficiency effect (detection_alpha) = 20. The training was stopped after 50,000 iterations. All other parameters were used at default settings. Cell2location estimates the posterior distribution of cell abundance of every cell type in every spot. Posterior distribution was summarised as 5% quantile, representing the value of cell abundance that the model has high confidence in, and thus incorporating the uncertainty in the estimate into values reported in the paper and used for downstream co-location analysis.

### Identification of cell-types located inside the thyroid

We classified specific cell types as contaminants from surrounding tissues if they were clearly biologically irrelevant (i.e. chondrocytes, myocytes), or only detected in a specific age group (i.e. epithelial cells). The spatial distribution of the remaining cell types was evaluated using cell2location output. Immune cells were classified as residing ‘inside’ the thyroid.

### RNAscope

Fresh frozen cryosections (10µm) were fixed in chilled 4% PFA for 15 mins and dehydrated through an ethanol series (50%, 70%, 100%, 100% ethanol, 5 mins each). Slides were processed on a Leica BOND RX system according to the RNAscope Multiplex Fluorescent Reagent Kit v2 Assay (Advanced Cell Diagnostics, Bio-Techne). Probes used are found in **Supplementary Table 5**. Prior to probe hybridisation sections underwent pre-treatment with Protease III, 15 mins at 37°C. Probe channels were developed using TSA-Opal dyes (Opal 520, Opal 570 and Opal 650; 1:1000; Akoya Biosciences), TSA-biotin (TSA Plus Biotin Kit, Perkin Elmer) and streptavidin-conjugated Atto 425 (1:300, Sigma Aldrich).

### H&E staining

Fresh frozen samples were embedded and frozen in OCT and sectioned using a Leica CM1950 cryostat. 10 µm cryosections were air dried, stained with Gills II Haematoxylin (Leica, 3801521E) for 1 min, followed by Aqueous Eosin Y (Sigma HT110216) for 3s, before dehydration (70% Ethanol, 100% ethanol), clearing (Neoclear) and coverslipping (Neomount). They were imaged using a Hamamatsu Nanozoomer 2.0 HT brightfield slide scanner system.

### Immunofluorescence

10 µm fresh frozen cryosections were fixed in 4% PFA for 20 min and then underwent automated staining on a Leica BOND RX system. Nonspecific antibody binding was blocked with 5% bovine serum albumin (BSA; Sigma Aldrich), 0.1% Tween-20 (BioRad) PBS solution for 1 hour at RT. Sections were then incubated with the primary antibody SLC5A5 (Proteintech, 24324-1-AP; 1:50), for 1 hour at RT, followed by incubation with the secondary antibody (Jackson 711-165-152; 1:500) for 1 hour at RT. Sections were stained with nuclear counterstain DAPI (Invitrogen, D1306; 1:50,000) for 30 mins at RT.

### Imaging

Fluorescently stained slides were imaged on an Opera Phenix system (Perkin Elmer), in confocal mode with 1µm z-step size, using a 63x water immersion lens. Channels were as follows: DAPI (excitation 375 nm, emission 435–480 nm), Atto 425 (excitation 425 nm, emission 463–501 nm), Opal 520 (excitation 488 nm, emission 500–550 nm), and Opal 570/Cy3 (excitation 561 nm, emission 570–630 nm).

#### Thyroid cancer analyses

##### Bulk RNA sequencing data acquisition and processing

###### TCGA data

Bulk transcriptomes (tables of raw counts) of adult thyroid tissues generated by the TCGA Research Network^54^, were obtained from the TCGA GDC data portal (https://portal.gdc.cancer.gov). 323 tumour samples with classical thyroid cancer diagnosis, along with 59 samples of normal thyroid tissue were included in the analysis (**Supplementary Table 7)**.

###### St Jude data

Bulk transcriptomes (tables of raw counts) of paediatric thyroid tumour samples, generated by St. Jude Children’s Research Hospital, were obtained from St. Jude Coud^53^. Overall, 37 relevant samples were included in the analysis (**Supplementary Table 7)**.

###### GTeX data

Bulk transcriptomes (tables of raw counts) of adult thyroid tissues from non-diseased sites were obtained from the Adult Genotype Tissue Expression (GTEx) project data portal (https://gtexportal.org/home/). Overall, 651 relevant samples were included in the analysis (**Supplementary Table 7)**.

Count tables were generated independently by the respective sources, all of which utilised the GRCh38 reference genome.

###### Yoo et al. 2016^90^

Raw sequencing data were retrieved from EBI European Nucleotide Archive database with accession number PRJEB11591. Using the STAR algorithm, sequencing reads were mapped to the GRCh38 reference genome and tables of counts were generated. 76 samples of classical papillary thyroid cancer in adults, along with 81 matched-normal thyroid tissues were included in the analysis (**Supplementary Table 7)**.

### Cell signal analysis comparing bulk transcriptomes to a single-cell RNAseq reference

With our annotated foetal thyroid scRNA-seq atlas as the reference, the transcriptional signal contribution of each cell type towards each bulk transcriptome was calculated using CellSignalAnalysis method as previously described in Young et.al 2021^55^. Signal contributions of endothelial cells, mesenchymal cells, myocytes, smooth muscle cells and ENS neurons were aggregated into the “Stromal (non-immune)” group (**Figure 4a**). Similarly, the “Immune” signal was the total signals across all immune cell types (B cells, monocytes, myeloid, NK and T cells); and the “Others” signal includes chondrocytes, glial cells and epithelial cells. As the method expects that no normal single-cell reference can perfectly match the cancer transcriptomes, any unexplained transcriptional signal is captured by the “Intercept/Unexplained” term. A one-tailed Wilcoxon test was performed to assess whether the fTFC2 signal is significantly more enriched in paediatric papillary thyroid cancer compared to adult papillary thyroid cancer (from TCGA and Yoo et al. 2016^90^).

### Whole genome sequencing

Whole genome sequencing data were generated for biopsies from primary tumour sites and normal thyroid tissue from both patients (**Supplementary Table 7**). DNA was extracted from frozen samples using the AllPrep DNA/RNA/Protein Mini Kit (QIAGEN) following the standard protocol. Short insert (∼500bp) genomic libraries were then constructed. Finally, 150-bp paired-end sequencing clusters were generated on the Illumina Novaseq 6000 platform according to Illumina standard library generation (with PCR) protocols. The average sequence coverage was 30X for both tumours and matched normal samples.

### Variant detection analyses

Raw DNA sequencing data were aligned to the GRCh38 (Ensembl 103) reference genome using the Burrows-Wheeler algorithm (BWA-MEM)^91^. For each patient, different classes of somatic variants were called using the well-established pipelines at the Wellcome Sanger Institute: single-nucleotide substitutions were called using CaVEMan algorithm (v1.18.2)^92^, short insertions/deletions were called using Pindel algorithm (v3.10.0)^93^, copy number variants were called with ASCAT (v4.5.0)^94^ and Battenberg (v3.5.3)^95^, and structural rearrangements were called using BRASS algorithm (v6.3.4)^96^.

### 10X single-nucleus RNA sequencing (snRNA-seq)

Single nuclei were isolated from frozen tissue using a glass dounce homogeniser. Samples were homogenised in buffer A (Sucrose 0.25 M, BSA 10 mg/ml, MgCl2 0.005 M, protease inhibitors and RNAse inhibitors RNAseIn 0.12 U/ul and Superasin 0.06 U/ul), using ∼25 strokes with the “loose” pestle and ∼20 strokes with the “tight” pestle. Nuclei were cleaned up using a 30% Percol gradient and resuspended in buffer B (Sucrose 0.32 M, BSA 10 mg/ml, CaCl2 3 mM, MgAc2 2 mM, EDTA 0.1 mM, Tris-HCl 10 mM, DTT 1 mM in the presence of protease and RNAse inhibitors as in buffer A). Nuclei were mixed 1:1 with Trypan blue and counted using a disposable haemocytometer, then diluted to the appropriate concentration. Nuclei were loaded onto the 10X Chromium controller as per the Chromium Next Gem Single Cell 5’ Kit V2 User Guide, targeting to recover 7000 nuclei. Post GEM-RT clean-up, cDNA amplification, and 5′ gene expression library construction were carried out according to the user guide. The resulting libraries were sequenced on the Novaseq 6000 platform.

### Quality control and preprocessing of snRNA-seq data

Raw sequencing data was processed and mapped to the GRCh38 2020-A human reference genome, using Cell Ranger pipeline (v7.1.0)^97^. The filtered count matrix, outputted by Cell Ranger, was then further QC’ed using Seurat (v4.0.1) in R (v4.0.4). Nuclei with <300 genes, <500 UMIs, or mitochondrial fraction exceeding 30% were removed. Scrublet (v0.2.3)^82^ was used to identify doublets. Nuclei were excluded if identified as doublets by Scrublet, or having a doublet score > 0.5. Ambient mRNA contamination was removed with SoupX (v1.6.1)^81^. High resolution Louvain clusters (resolution=10) with >50% cells failing QC were also excluded. Data were log normalised and scaled, and principal components were calculated using the top 2000 highly variable genes, following the standard Seurat workflow. Louvain clustering was performed (resolution = 1), and a uniform manifold approximation and projection (UMAP) calculated, using the top 75 principal components. No integration or batch correction method was performed to preserve the biological variation across different tumour samples.

### Annotation of snRNA-seq paediatric thyroid dataset

A semi-automated cell type annotation by label transfer approach was employed. Using Celltypist^56^, a logistic regression model was trained on the reference scRNA-seq foetal thyroid atlas. The model is then used to calculate the predicted similarity scores of individual cells in our query snRNA-seq dataset against each reference class (i.e. different cell types in the reference dataset). Cells are assigned the label of the class with the highest positive similarity score. Annotation was further refined by manual assessment of well-established cell-type specific marker genes in high-resolution clusters (**Supplementary Figure 7e**). Tumour cells were identified by the loss of *TPO* whilst maintaining the expression of other thyrocyte markers, including *TSHR*, *PAX8*, and *TG*.

### fTFC1 and fTFC2 gene signature in sc/snRNA-seq data

Genes were included in the fTFC1 and fTFC2 gene signature based on the following criteria:

1. identified as a significantly differentially up-regulated in fTFC1 or fTFC2 respectively, as defined in the analysis described above;
2. expressed in at least 50% fTFC1 or fTFC2 cells, respectively;
3. expressed in at least 30% normal thyrocytes in the snRNA-seq dataset.

To prioritise genes which are highly specific to each population, the selected genes were then ranked by the difference in percentage of fTFC1 and fTFC2 cells with positive expression. Top 70 genes from each list were included in the gene signatures. Normal thyrocytes and papillary thyroid cancer cells were extracted from our scRNA-seq foetal atlas and snRNA-seq paediatric dataset, as well as published adult scRNA-seq datasets^24,45,98^. The enrichment score of each gene signature in individual cells / nuclei was calculated using the function ‘AddModuleScore_UCell’ from the R package UCell (v1.3.1)^99^.

## Data availability

High-throughput raw sequencing data in this study is available upon request. Bulk transcriptomes of paediatric papillary thyroid cancer used for analysis in this study were obtained from St. Jude Cloud^53^ (https://www.stjude.cloud). We also utilised publicly available data of adult normal thyroid tissue and adult papillary thyroid cancer from the following sources: (1) bulk transcriptomes were obtained from the TCGA GDC data portal (https://portal.gdc.cancer.gov), the Genotype-Tissue Expression (GTEx) project^100^ (https://gtexportal.org/home), and Yoo et al 2016^90^; and (2) single-cell RNAseq data were obtained from Mosteiro et al. 2023^24^, Wang et al. 2022^45^, and Pu et al. 2021^98^.

## Acknowledgments

This publication is part of the Human Cell Atlas – www.humancellatlas.org/publications/. The authors thank the Sanger Cellular Generation and Phenotyping (CGaP) Core Facility and the Sanger Core Sequencing pipeline for support with sample processing and sequencing library preparation. Thyroid developmental material was provided by the Joint MRC–Human Cell Atlas (MR/S036350/1). The authors are grateful to patients for donating tissue for research; Luz Garcia-Alonso, Valentina Lorenzi, Kevin Troulé, Nathaniel Anderson and Matthew Young for bioinformatics advice; Loren Gibson for ethics and contracts support; Tarryn Porter and Heather Stanley for logistics support, Cellular Genetics wet lab team for experimental support; Bee Ling Ng and the Cytometry Core Facility for cell sorting support; Kenny Roberts for microscopy advice; Human Developmental Biology Resource (HDBR) for tissue coordination support; Antonio García from Bio-Graphics for scientific illustrations; Aidan Maartens for proofreading.

## Author information

H.M, M.K.T, S.B, R.V-T and N.S conceived and designed the experiments and analyses. H.M, and M.K.T analysed the data with contributions from E.A, A.P, P.M, H.J.W. H.M performed sample processing with help from C.S-S, A.O, Y.W, C.P, T.O. L.T performed the imaging experiments. H.M, M.K.T, I.K, S.B, R.V-T and N.S interpreted the data. H.M, M.K.T, S.B, R.V-T and N.S wrote the manuscript. N.S, R.V-T and S.B supervised the work. All authors read and approved the manuscript.

